# Integration of multi-omics data reveals interplay between brassinosteroid and TORC signaling in Arabidopsis

**DOI:** 10.1101/2022.03.11.484017

**Authors:** Christian Montes, Ping Wang, Ching-Yi Liao, Trevor M Nolan, Gaoyuan Song, Natalie M Clark, J. Mitch Elmore, Hongqing Guo, Diane C Bassham, Yanhai Yin, Justin W Walley

## Abstract

Brassinosteroids (BR) and Target of Rapamycin Complex (TORC) are two major actors coordinating plant growth and stress responses. BRs function through a signaling pathway to extensively regulate gene expression and TORC is known to regulate translation and autophagy. Recent studies revealed that these two pathways crosstalk, but a system-wide view of their interplay is still missing. Thus, we quantified the level of 23,975 transcripts, 11,183 proteins, and 27,887 phosphorylation sites in wild-type Arabidopsis and in mutants with altered levels of either BRASSINOSTEROID INSENSITIVE 2 (BIN2) or REGULATORY ASSOCIATED PROTEIN OF TOR 1B (RAPTOR1B), two key players in BR and TORC signaling, respectively. We found that perturbation of BIN2 or RAPTOR1B levels affects a common set of gene-products involved in growth and stress responses. Furthermore, we used the multi-omic data to reconstruct an integrated signaling network. We screened 41 candidate genes identified from the reconstructed network and found that loss of function mutants of many of these proteins led to an altered BR response and/or modulated autophagy activity. Altogether, these results establish a predictive network that defines different layers of molecular interactions between BR-or TORC-regulated growth and autophagy.

## Introduction

Organisms are frequently affected by environmental challenges. When responding to stress, specific molecular and cellular processes are triggered, and growth is often compromised. These responses to both biotic and abiotic stresses rely heavily on modulating hormonal signaling pathways, and plants need to allocate resources between their growth and stress response machinery efficiently. Therefore, well-coordinated hormonal crosstalk is fundamental for a successful response to stress (Huot et al., 2014; Verma et al., 2016; Bürger and Chory, 2019). The growth-promoting hormone brassinosteroid (BR) has been shown as a critical element in this balance. Plants with altered levels of BR signaling or biosynthesis genes exhibit deficient growth (Li et al., 1996; Li and Chory, 1997; Li and Nam, 2002; Yin et al., 2002; Chung et al., 2010; Guo et al., 2013) and abnormal response to various stresses (Che et al., 2010; Nolan et al., 2017b; Ye et al., 2017; Fàbregas et al., 2018; Gruszka, 2018; Planas-Riverola et al., 2019; Xie et al., 2019; Gupta et al., 2020; Liang et al., 2020).

The GLYCOGEN SYNTHASE KINASE 3 (GSK3)-like kinase BRASSINOSTEROID INSENSITIVE 2 (BIN2) is a critical negative regulator of BR signaling (Li and Nam, 2002; Kim et al., 2009). In the absence of BRs, BIN2 phosphorylates the bri1-EMS-SUPPRESSOR1/BRASSINAZOLE RESISTANT1 (BES1/BZR1) family of transcription factors (TFs), which reduces their protein level, lowers DNA binding, and promotes cytoplasmic sequestration by 14-3-3 proteins, thereby preventing the activation of downstream BR response genes (Yin et al., 2002; Gampala et al., 2007; Ryu et al., 2007, 2010). BR signals through the receptor BRI1, coreceptor BAK1, and other components to inhibit BIN2, allowing BES1/BZR1 accumulation in the nucleus to regulate thousands of BR genes for various BR responses (Li and Chory, 1997; Wang et al., 2001; Nam and Li, 2002; Kim et al., 2009; Zhu et al., 2017; Nolan et al., 2020). Besides regulating BES1 and BZR1, increasing evidence position BIN2 as a hub for regulation of the balance between stress and growth (Youn and Kim, 2015; Nolan et al., 2020). BIN2 is involved in BR-regulation of diverse processes such as drought and abscisic acid (ABA) signaling (Cai et al., 2014; Hu and Yu, 2014; Ye et al., 2017; Jiang et al., 2019), cold stress response (Ye et al., 2019), salt-stress response (Li et al., 2020a), root development in conjunction with auxin signaling (Cho et al., 2014; Li et al., 2020b) as well as chloroplast development (Zhang et al., 2021). Despite the increasing number of reports with BIN2 acting as an essential regulator in growth/stress balance, no multi-omics studies on this kinase have been reported so far.

In *Arabidopsis thaliana*, the TARGET OF RAPAMYCIN complex (TORC) is an important regulator that integrates nutrient and energy sensing into cell proliferation and growth (Xiong and Sheen, 2014; Fu et al., 2020). Activation of TORC signaling induces the expression of ribosomal proteins, increases protein translation, stimulates photosynthesis, and upregulates (transcriptionally and translationally) plant growth-promoting genes (Ren et al., 2012; Xiong et al., 2013; Dong et al., 2015; Van Leene et al., 2019; Scarpin et al., 2020). Conversely, TORC actively represses autophagy, a central recycling system of cytoplasmic components that is essential for rerouting nutrients and other raw materials when needed for plant growth, development, or stress responses (Noda and Ohsumi, 1998; Pu et al., 2017; Marshall and Vierstra, 2018). TORC is comprised of TOR kinase, LETHAL WITH SEC THIRTEEN PROTEIN 8 (LST8), and REGULATORY ASSOCIATED PROTEIN OF TOR (RAPTOR). TOR is the catalytic component of TORC, LST8 provides stability, and RAPTOR interacts with and recruits substrates to the complex (Hara et al., 2002; Mahfouz et al., 2006; Yang et al., 2013). In Arabidopsis, null mutants in *TOR* are embryo lethal (Menand et al., 2002). Two RAPTOR homologs, RAPTOR1A and RAPTOR1B, have been found in Arabidopsis with *RAPTOR1B* being the predominantly expressed copy (Anderson et al., 2005; Deprost et al., 2005). *Raptor1a raptor1b* double mutant plants maintain embryonic development, unlike *tor* mutants, but lack post-embryonic growth (Anderson et al., 2005). *Raptor1a* null mutants have no major developmental phenotypes. Loss of function *raptor1b* mutants have reduced TORC activity (Wang et al., 2018; Salem et al., 2018). Consistently, *raptor1b* mutants exhibit reduced growth and development as well as increased basal levels of autophagy (Anderson et al., 2005; Salem et al., 2018; Pu et al., 2017).

When plants encounter stress, autophagy is often triggered, and growth-promoting pathways such as BR or TORC signaling need to be dampened (Nolan et al., 2017a; Liao and Bassham, 2020). To enable this balanced regulation of plant growth and stress responses, hormonal pathways such as auxin (Li et al., 2017; Schepetilnikov et al., 2017) and BRs (Zhang et al., 2016; Vleesschauwer et al., 2018) can influence or be affected by TORC activity. Increasing evidence points towards TORC-regulated autophagy as a crucial interaction point between BRs and TORC signaling when controlling this balance. For example, activation of TORC signaling promotes BR response by stabilizing BZR1, likely preventing its autophagy-driven degradation (Zhang et al., 2016). Additionally, BIN2 knock-down lines exhibit reduced sensitivity to TOR inhibitors AZD8055 (AZD) and KU63794 (Xiong et al., 2017). Furthermore, RIBOSOMAL PROTEIN S6 KINASE 2 (S6K2) can phosphorylate BIN2 in a TOR-dependent manner. However, the mechanism and biological implications of this interaction are not clear (Xiong et al., 2017). Under stress conditions such as drought or sucrose starvation, BES1 is ubiquitinated by SINAT2 and/or BAF1 ubiquitin ligases and targeted to selective autophagy through ubiquitin receptor DSK2 to slow down plant growth (Nolan et al., 2017b; Yang et al., 2017; Wang et al., 2021). Moreover, BIN2 has been shown to phosphorylate ubiquitin receptor DSK2 to facilitate its interaction with ATG8 and promote BES1 degradation via selective autophagy (Nolan et al., 2017b).

BIN2 and TORC regulate plant responses to environmental changes via phosphorylation, exerting molecular changes at many different levels (i.e., changes in gene transcription or protein activity) (Guo et al., 2013; Youn and Kim, 2015; Bozhkov, 2018; Van Leene et al., 2019; Liao and Bassham, 2020; Nolan et al., 2020). Therefore, understanding the molecular connection between BR and TORC signaling across different levels of gene expression is necessary to unravel the interplay between these pathways. Furthermore, despite BIN2 being intensively studied, proteome-wide identification of BIN2 substrates is lacking. Here, we present a comprehensive multi-omic profiling detailing transcriptome, proteome, and phosphoproteome changes that occur in mutants with altered levels of BIN2 or the TORC subunit RAPTOR1B. We complement these global *in vivo* profiles with proteome-wide identification of direct BIN2 substrates using a Multiplexed Assay for Kinase Specificity (MAKS). Substantial overlap was found in the transcripts, proteins, and phosphosites whose accumulation is dependent on BIN2 and RAPTOR1B. Using this wealth of information, we reconstructed an integrated kinase-signaling network and gene regulatory network (GRN). We used this integrated network to identify novel genes whose mutant lines showed either altered growth in response to BR and/or levels of autophagy. Together, these studies further our understanding of the dynamic interplay between BR and TORC signaling.

## Results

### Multi-omics profiling of *bin2* and *raptor1b* mutants provides insights into known and new regulatory roles

We designed a multi-omics experiment to identify novel components dependent on BR and/or TORC signaling. We performed transcriptome, proteome, and phosphoproteome profiling on rosette leaves of 20-day old wild-type (WT), *bin2D* (gain-of-function), *bin2T* (*bin2 bil1 bil2* triple loss of function), and *raptor1b* loss of function plants. We quantified transcript levels using 3’ QuantSeq (Moll et al., 2014) and measured protein abundance and phosphorylation state using two-dimensional liquid chromatography-tandem mass spectrometry (2D-LC-MS/MS) on Tandem Mass Tag (TMT) labeled peptides (McAlister et al., 2012; Hogrebe et al., 2018; Song et al., 2018a) (Figure 1A). From these samples, we detected 23,975 transcripts, 11,183 proteins, and up to 27,887 phosphosites from 5,675 phosphoproteins (Figure 1B and Supplemental Data Set S1). We found 5,653 transcripts and 4,001 protein groups (hereafter referred to as proteins) that were differentially expressed (DE) in at least one mutant when compared to WT (Figure 1C and Supplemental Figure S1, A and B). Gene ontology (GO) analysis of DE transcripts and proteins in *bin2D, bin2T*, and *raptor1b* mutants showed enrichment of many terms from similar processes including growth, hormones, stimuli sensing, and stress (Figure 2 A and B; Supplemental Data Set S2 and S3), which is consistent with the known roles of BIN2 and RAPTOR in growth/stress balance and hormonal crosstalk.

**Figure 1.**
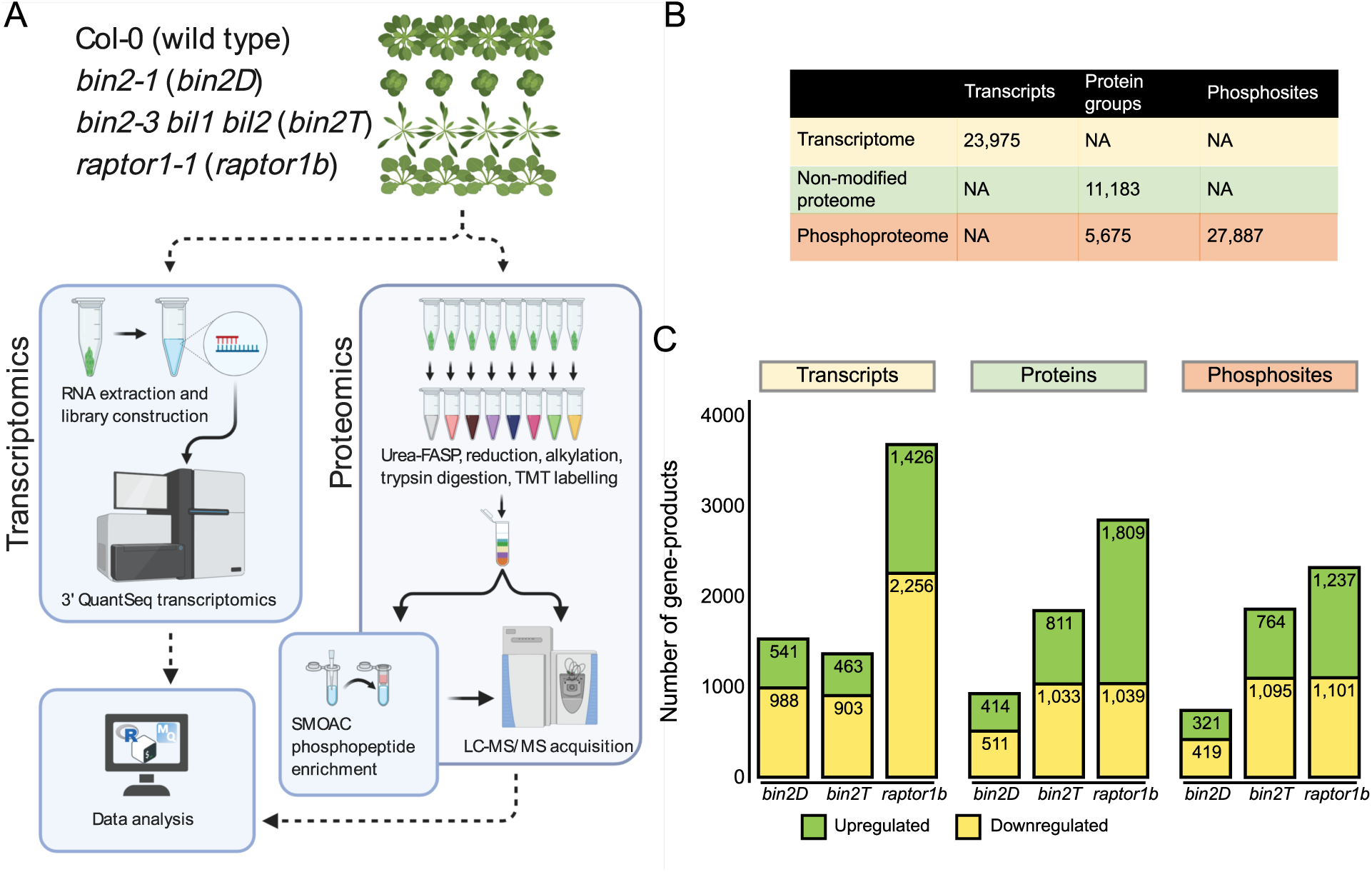
Experimental design, workflow, and data overview. **A**, Schematic representation of the multi-omics processing pipeline for *bin2* and *raptor1b* mutants. **B**, number of total detected transcripts, proteins, phosphoproteins, or phosphosites. **C**, differentially expressed transcripts, proteins, and phosphorylated amino acids for each analyzed mutant compared to WT.

**Figure 2.**
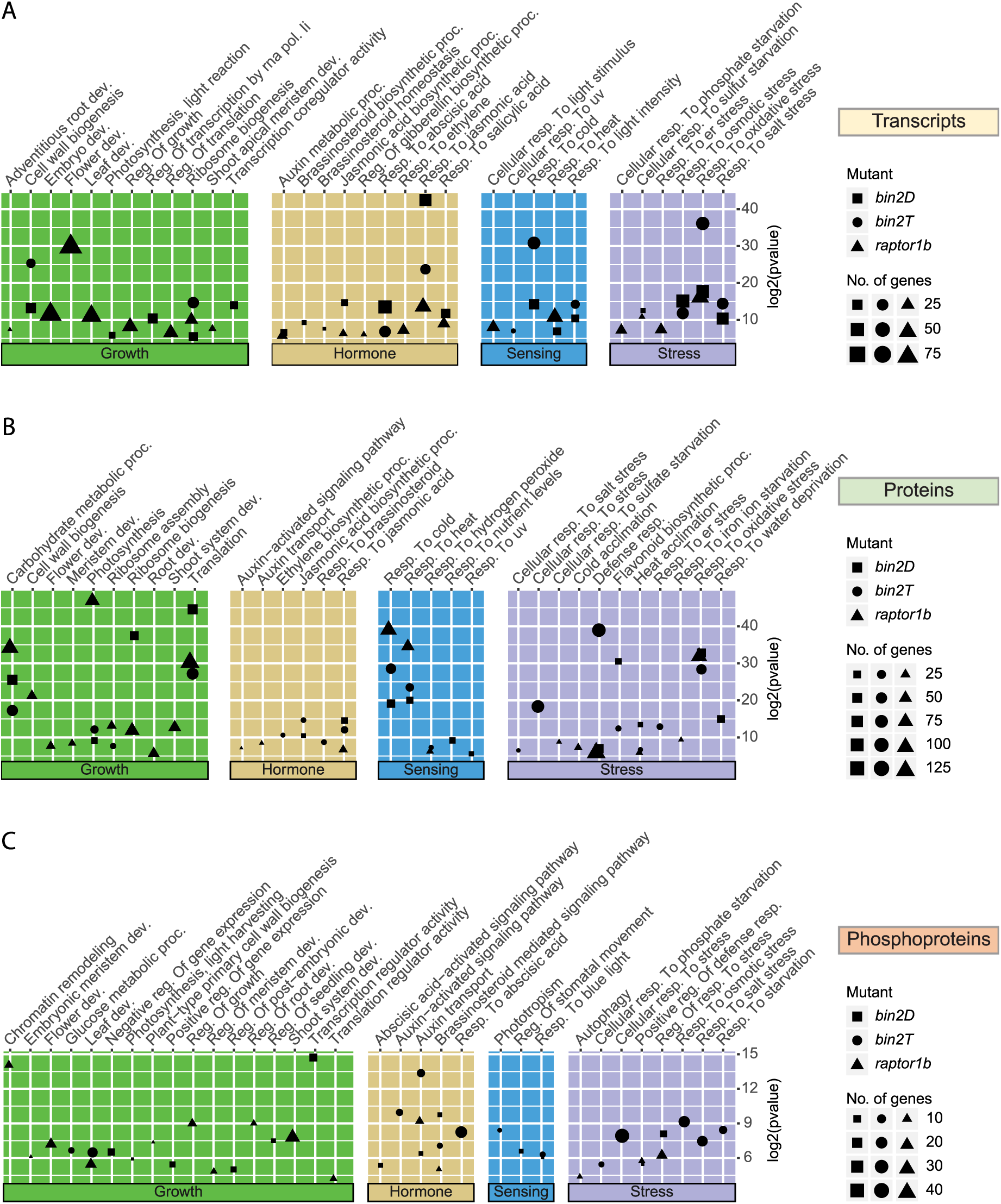
Gene ontology analysis on datasets. Selection of significant GO biological processes among differential expressed transcripts (**A**), proteins (**B**), and phosphosites (**C**), on *bin2D* (■), *bin2T* (●), and *raptor1b* (▲). For transcripts and protein expression, significant terms were selected from both, up-and downregulated genes. For phosphosites, terms were selected from those differentially expressed in the directionality of the respective kinase mutant (i.e., upregulated in *bin2D*, or downregulated in *bin2T* or *raptor1b*)

Our phosphoproteomics analysis identified 4,153 differentially phosphorylated sites in at least one mutant (Figure 1C, Supplemental Data Set 1C, and Supplemental Figure S1C). GO analysis for potential BIN2 target proteins (i.e., those with increased phosphorylation in *bin2D* or decreased phosphorylation in *bin2T*) revealed enrichment of terms related to plant growth and development, as well as response to stress and defense, processes in accordance with known functions of BIN2 and its homologs. In addition, response to BR, abscisic acid, and auxin terms were also significant, highlighting once more the close relationship between BIN2 activity and these hormones. Transcriptional regulation-related terms were significantly enriched in the *bin2D* dataset, consistent with BIN2’s well documented regulatory activity upon transcription factors (TFs) (Figure 2C and Supplemental Data Set S4). Finally, we assessed GO enrichment for proteins with decreased phosphorylation in *raptor1b*. We found that most of the enriched terms were related to growth, autophagy, starvation, auxin, and BR response. This is consistent with the known biological role of RAPTOR and suggests a cross-regulation between BR and TORC pathways via phospho-signaling (Figure 2C and Supplemental Data Set S4).

### Phosphoproteomic analysis of *bin2* mutants shows enrichment of BIN2 direct targets

Because BIN2 is a kinase, we hypothesized that phosphosites increased in the *bin2D* gain-of-function mutant or decreased in *bin2T* may be direct BIN2 substrates. To test this hypothesis, we generated a proteome-wide dataset of BIN2 direct targets using the multiplexed assay for kinase specificity (MAKS) (Brumbaugh et al., 2014; Jayaraman et al., 2017) (See Methods for details). We quantified a total of 10,375 phosphosites accounting for 3,628 phosphoproteins from this assay (Supplemental Data Set S1C). As expected, the obtained phosphoproteome was heavily skewed toward increased phosphosites, with 1,343 phosphosites increasing following incubation with GST-BIN2 (Figure 3A and Supplemental Data Set S1C). Among proteins with increased phosphorylation we observed YDA and BSK1, two known BIN2 targets (Kim et al., 2012; Sreeramulu et al., 2013). To evaluate this set of phosphorylation sites as BIN2 kinase-substrates, we performed motif enrichment analysis and found a significant enrichment of the well-known GSK3 motif “S/T-X-X-X-S/T” (Fiol et al., 1987; Youn and Kim, 2015) among the increased phosphosites (p < 0.01, Figure 3B and Supplemental Data Set S5A). Additionally, another highly enriched motif found in the analysis was “S/T-P”, which is reported as a motif for GSK3, CDK, and MAPK families (Amanchy et al., 2007; Lin et al., 2015) (Supplemental Data Set S5A). Some previously unreported length variations of the GSK3 motif were also significantly enriched (i.e., S/T-X-X-S/T, S/T-X-S/T, and S/T-S/T, Supplemental Data Set S5A). These results support the robustness of our BIN2 kinase dataset and suggest a more flexible substrate recognition motif for BIN2 as a GSK3-like kinase.

**Figure 3.**
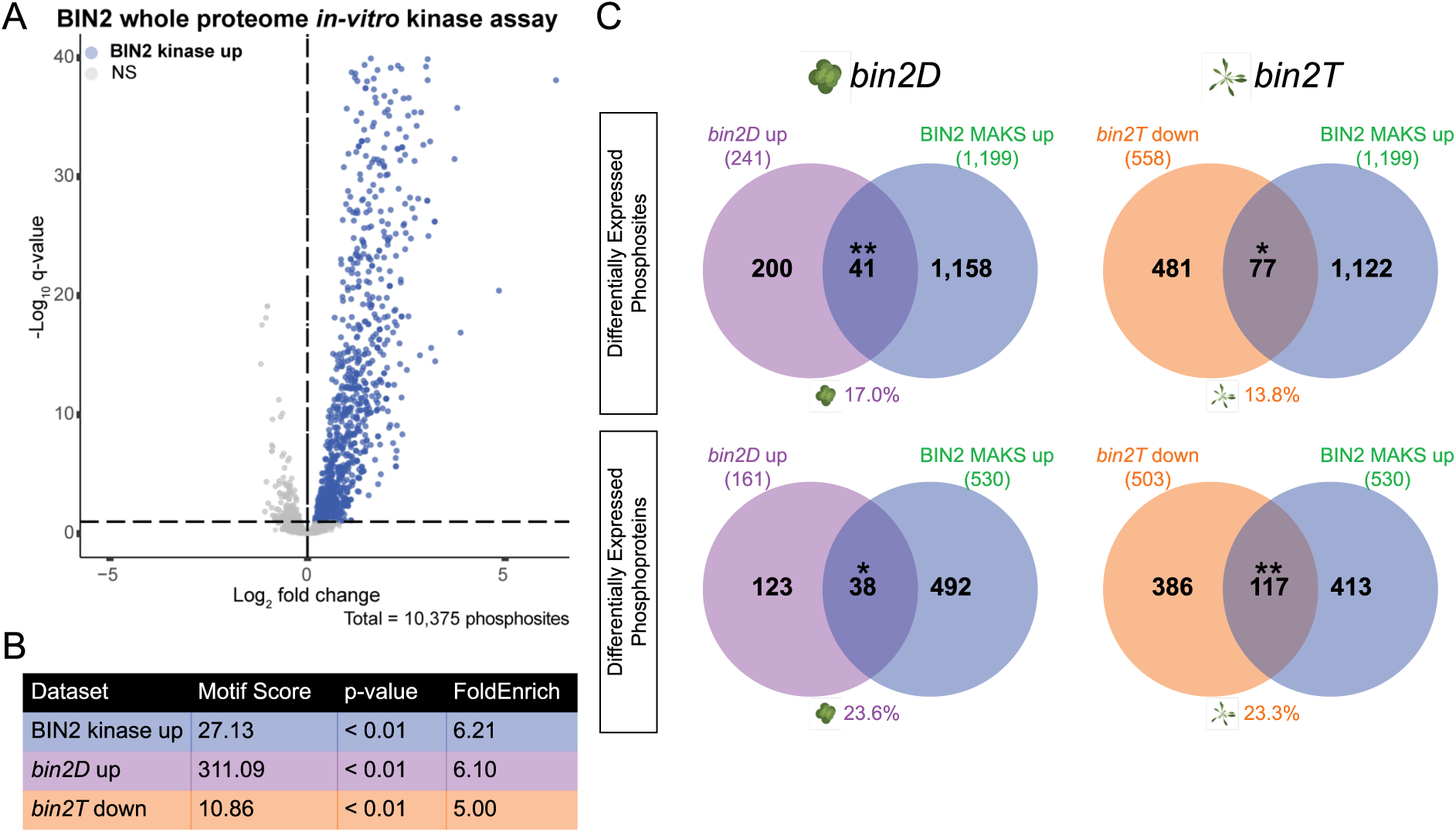
Phosphoproteomic analysis on *bin2* mutants shows significant enrichment of BIN2 direct targets. **A**, Volcano plot of phosphorylation sites from a Multiplexed Assay for Kinase Specificity (MAKS) on Arabidopsis leaf protein extracts incubated with recombinant GST or GST-BIN2. Significantly increased phosphosites are colored blue (q < 0.1). **B**, De novo motif enrichment analysis showed high enrichment for the GSK3 motif on BIN2-related phosphoproteomic datasets. Motif score and FoldEnrich values are calculated by motifeR, while *p*-value was calculated using hypergeometric testing. **C**, Venn diagrams show overlap between BIN2 direct targets (i.e., those upregulated in BIN2 MAKS) and phosphosites (upper) or phosphoproteins (lower) upregulated in *bin2D* (left) or downregulated in *bin2T* (right) mutants. Numbers below each Venn diagram represent the overlapping percent for that mutant (purple = *bin2D*, orange = *bin2T*). Phosphosite overlaps were calculated using a 20 amino acid window, centered on the differentially regulated phosphosite (for details see methods section). Statistical significance was calculated using hypergeometric testing (* *p* <0.05, ** *p* < 0.01).

We next assessed the prevalence of BIN2 direct targets present in our *in vivo* profiling of *bin2D* and *bin2T* mutants. For this, we first performed motif enrichment analysis on phosphosites perturbed in the expected direction for BIN2 targets (i.e., either upregulated in *bin2D* or downregulated in *bin2T*). A significant enrichment for the GSK3 motif was found in both *bin2D* up (p < 0.01) and *bin2T* down (p < 0.01) phosphosites (Figure 3B and Supplemental Data Set S5, B and C). Next, we looked at the overlap with BIN2 direct targets identified in the MAKS experiment. For *bin2D*, 17.0% (41/241; p < 0.01) of the total differentially upregulated phosphosites and 23.6% (38/161; p < 0.01) of the upregulated phosphoproteins were also BIN2 direct targets. For *bin2T*, 13.8% (77/558; p < 0.05) of the downregulated phosphosites and 23.3% (117/503; p < 0.01) of the downregulated phosphoproteins were also part of our BIN2 direct substrate list (Figure 3C and Supplemental Data Set S6). These results indicate that a subset of the BIN2-dependent phosphosites identified by *in vivo* mutant profiling may be direct BIN2 substrates.

### Kinase-signaling network inference on *bin2* and *raptor1b* mutants

Since both BIN2 and RAPTOR1B (TORC) participate in phosphorylation-based signaling, we reconstructed the molecular relationships of these signaling networks. To do so, we used our data to infer a kinase-signaling network for each mutant (i.e., *bin2D, bin2T*, and *raptor1b*). To build these networks, we inferred the activation state of kinases in our dataset. The activation loop (A-loop) is a well-conserved region inside the kinase domain whose phosphorylation is necessary for kinase activation (Adams, 2003; Ahiri, 2019). Thus, phosphosite intensity level of the A-loop can be used as a proxy for kinase activity quantification (Walley et al., 2013; Beekhof et al., 2019; Schmidlin et al., 2019; Clark et al., 2021). First, we performed a whole-proteome Arabidopsis *in-silico* A-loop prediction and were able to identify this region on 1,360 proteins (Supplemental Data Set S7A). Subsequently, we identified kinases whose A-loop phosphosite intensity was differentially regulated in at least one of the profiled mutants (Supplemental Figure S2A and Supplemental Figure S7B).

We found 27, 21, and 24 kinases exhibiting an altered activation state in the *bin2D, bin2T*, and *raptor1b* mutants, respectively (Supplemental Data Set S7B). Using this information, we inferred a kinase-signaling network by correlating phosphosite level with kinase activation state (Supplemental Figure S2B). A network containing 4,138 nodes, representing 33 activated kinases and 2,284 target sites arising from 1,853 possible substrate proteins was obtained (Figure 4A and Supplemental Data Set S8). To evaluate this kinase-signaling network, predicted BIN2 targets were obtained (i.e., nodes connected by edges directed outward of BIN2), and motif enrichment analysis was performed. As expected, the GSK3 motif was enriched among BIN2 targets (p = 1.96e-06). Additionally, the MAPK consensus motif P-X-[pS/pT]-P was overrepresented among MPK6 targets (p < 0.01). Several variants of the proline-directed phosphorylation motif [pS/pT]-P were significantly enriched among targets of MPK4 (p < 0.001), MPK10 (p < 0.001), and BIN2 (p < 0.001) (Amanchy et al., 2007; Lin et al., 2015; Rayapuram et al., 2021) (Figure 4B and Supplemental Data Set S9). Finally, 70% (7/10) of known BIN2 targets reported in the literature and present in our network were correctly predicted as BIN2 targets (either as direct targets or direct downstream second neighbor, Supplemental Data Set S8B). These results support the target prediction value of our inferred kinase signaling network.

**Figure 4.**
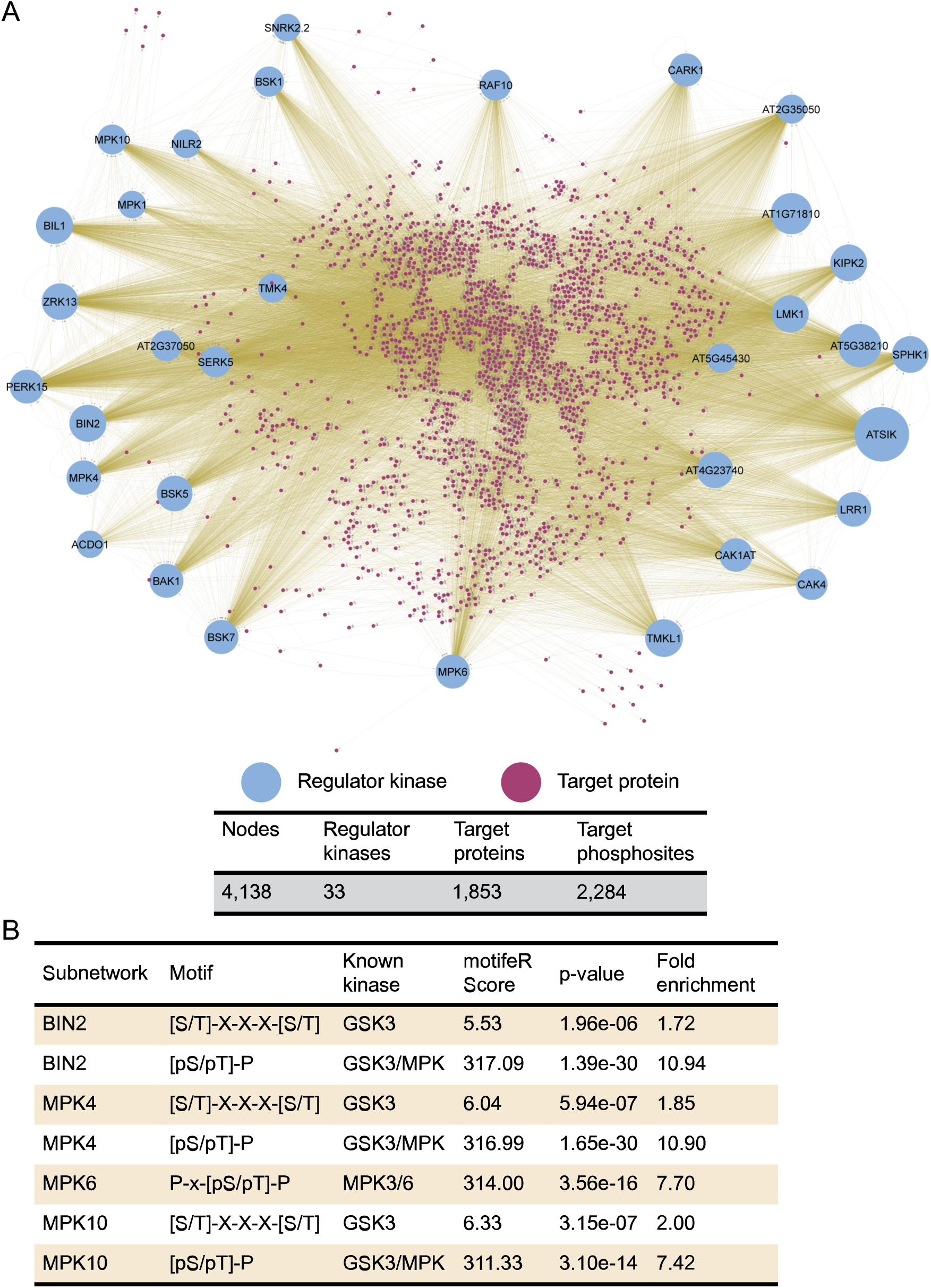
Kinase signaling network. **A**, A signaling network was inferred using phosphoproteomic data from *bin2D, bin2T*, and *raptor1b* mutants. Activated kinases are shown as named circles with their size representing the number of predicted targets (i.e., node outdegree). Target proteins are represented as small, purple circles. **B**, De novo motif enrichment analysis among predicted direct targets (i.e., node first neighbors) for BIN2, MPK4, and MPK10 showed high enrichment for the GSK3 motif ([S/T]-X-X-X-[S/T]) and GSK3/MPK3 motif ([pS/pT]-P). Analysis on MPK6 predicted direct targets showed significant enrichment of MPK3/6 motif (P-X-[pS/pT]-P). Enrichment analysis was done on a 14 amino acids window, centered on target phoshposites.

### Integrative multi-dimensional signaling network reconstruction reveals proteins required for normal BR response and autophagy

We have previously shown that using multiple omics datasets (i.e. transcriptomic, proteomic, and phosphoproteomic data) can increase the predictive power of Gene Regulatory Network (GRN) inference (Walley et al., 2016). With this in mind, we also inferred two separate transcription factor (TF)-centered GRNs, for each of our mutants (*bin2D, bin2T*, and *raptor1b*), using the SC-ION pipeline (Figure 5) (Clark et al., 2021). In the first network, called “abundance GRN”, TF protein abundance (when quantified) or TF transcript abundance (when cognate protein was not quantified) was used as the “regulator” value to infer their “target” transcript abundance. (Figure 5, blue line). The second network, termed “phosphosite GRN”, uses the TF phosphorylation intensity value as “regulator” to predict “target” transcript abundance (Figure 5, green line). We then integrated these two GRNs with the kinase-signaling network to provide a, multi-layered, portrait of signaling cascades (Figure 5; Supplemental Data Set S10). In this network, there are 2,272 BIN2-responsive nodes (kinases, regulatory TFs, or targets) that are present based on regulatory inference being made using information from mutants with altered BIN2 levels (*bin2D, bin2T*, Figure 5, left side). At the same time, there are 2,370 nodes only present based on the *raptor1b* mutant that can be thought of as primarily RAPTOR1B/TORC-responsive elements (Figure 5, right side). Additionally, 1,044 nodes are present based on inference made due to altered BIN2 and RAPTOR1B expression, showing crosstalk between both signaling cascades (Figure 5, center). To further characterize this integrated network, and to identify important regulators, we calculated the Integrated Value of Influence (IVI) score for each node, which integrates network centrality measurements into one normalized value to account for each node’s ranked importance in the analyzed network (Salavaty et al., 2020). Among the top 10% most influential nodes, we observe well-known BR signaling regulators such as BAK1, BEH1, BES1, BIM1, BIN2, and BSK1, as well as previously reported TOR targets such as VIP1, RBR1, and HAG1 (Van Leene et al., 2019) (Supplemental Data Set S10B).

**Figure 5.**
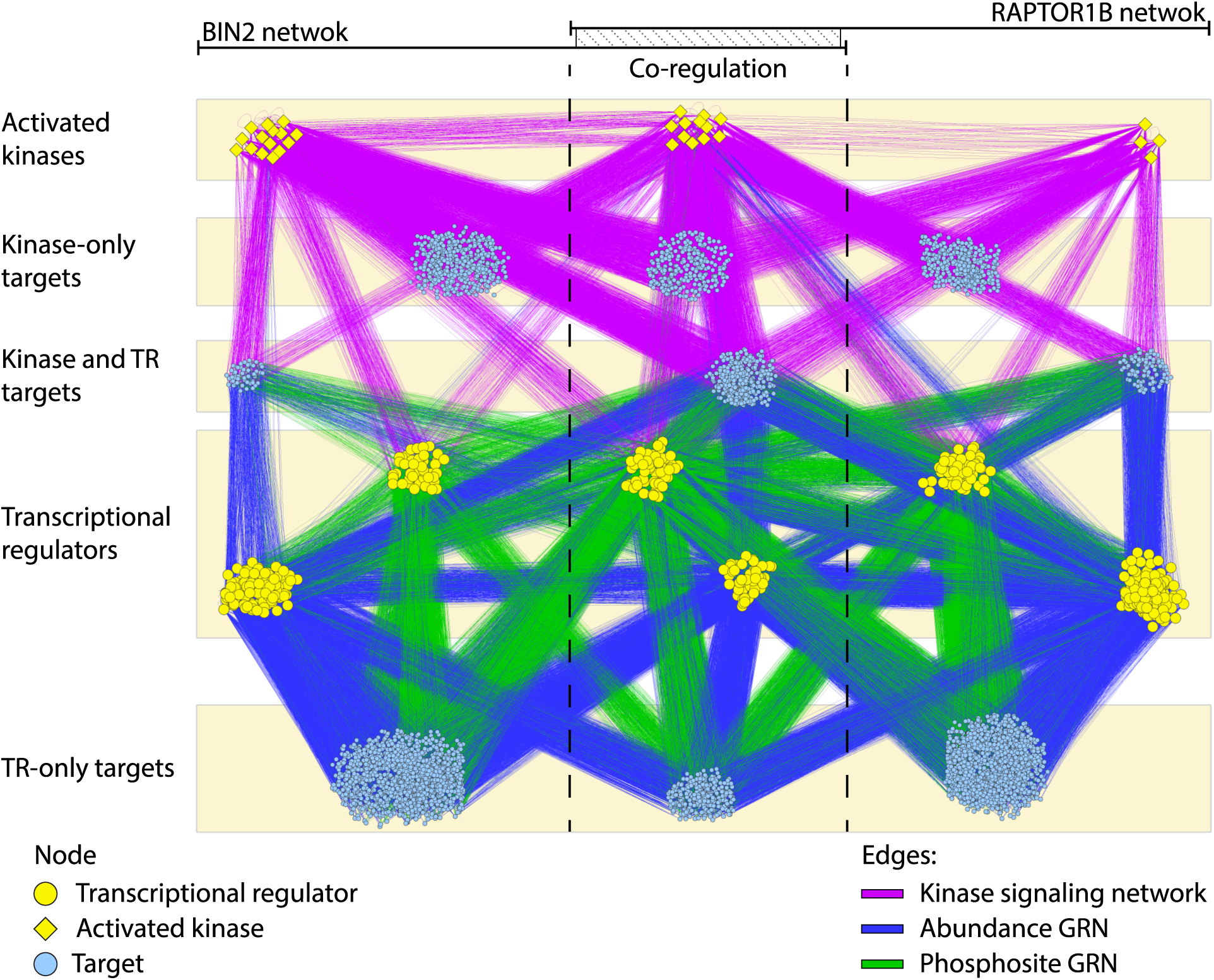
Multi-dimensional integrative network. Kinase-signaling network (purple lines), abundance GRN (blue lines), and phosphosite GRN (green lines) were reconstructed for each mutant (i.e., *bin2D, bin2T*, and *raptor1b*) using transcriptomic, proteomic and phosphoproteomic information and merged into an integrative network (see methods).

Next, we examined the integrative signaling network to determine whether these predictions identified proteins involved in BR response and/or TORC-response. To assess this question, we enlisted those proteins being differentially phosphorylated simultaneously in either of the BIN2 mutants (i.e., *bin2D* or *bin2T*) and in *raptor1b* (Figure 6). When selecting candidates, we focused our attention first on those proteins up-phosphorylated in *bin2D* and those down-phosphorylated in *bin2T* since this phosphorylation “directionality” could pinpoint those proteins direct or indirectly affected by BIN2 activity. We then fine-tuned this selection into possible BIN2/RAPTOR1B crosstalk by keeping only those proteins exhibiting differential phosphorylation in *raptor1b* (Figure 6). From this universe, we selected 41 candidate genes that are also present in our integrative network as crosstalk elements (Figure 5, center) and obtained mutants to assess their BR and autophagy phenotypes (Supplemental Data Set S11). Mutant lines for these candidate genes were identified and tested for hypocotyl elongation in response to BL as a means to assess their sensitivity to BR. From the tested candidate genes, 31.7% (13/41) showed a significantly altered BL response (Figure 7, and Supplemental Data Set S11A). We next measured autophagy levels as a readout of TORC activity, a total of 29 candidate genes that showed significantly altered hypocotyl elongation in response to BL and/or exhibited decreased phosphorylation in *raptor1b* mutant were examined for autophagy activity by transient expression of a GFP-ATG8e marker, which labels autophagosomal membranes, in protoplasts obtained from mutant lines (Contento et al., 2005). Twenty genes (69% of assayed candidate genes) showed significantly altered basal autophagy levels when mutated, with 15 of them being higher than WT and five lower than WT (Figure 8 and Supplemental Data Set S11B). GFP-ATG8e was also assessed under sucrose starvation for the same genotypes. Nineteen genes (65.5% of assayed genes) showed significant changes in autophagosome number under sucrose starvation conditions. Interestingly, there was little to no increase in autophagy upon sucrose starvation in the five genotypes with low basal autophagy (Figure 8 and Supplemental Data Set S11B).

**Figure 6.**
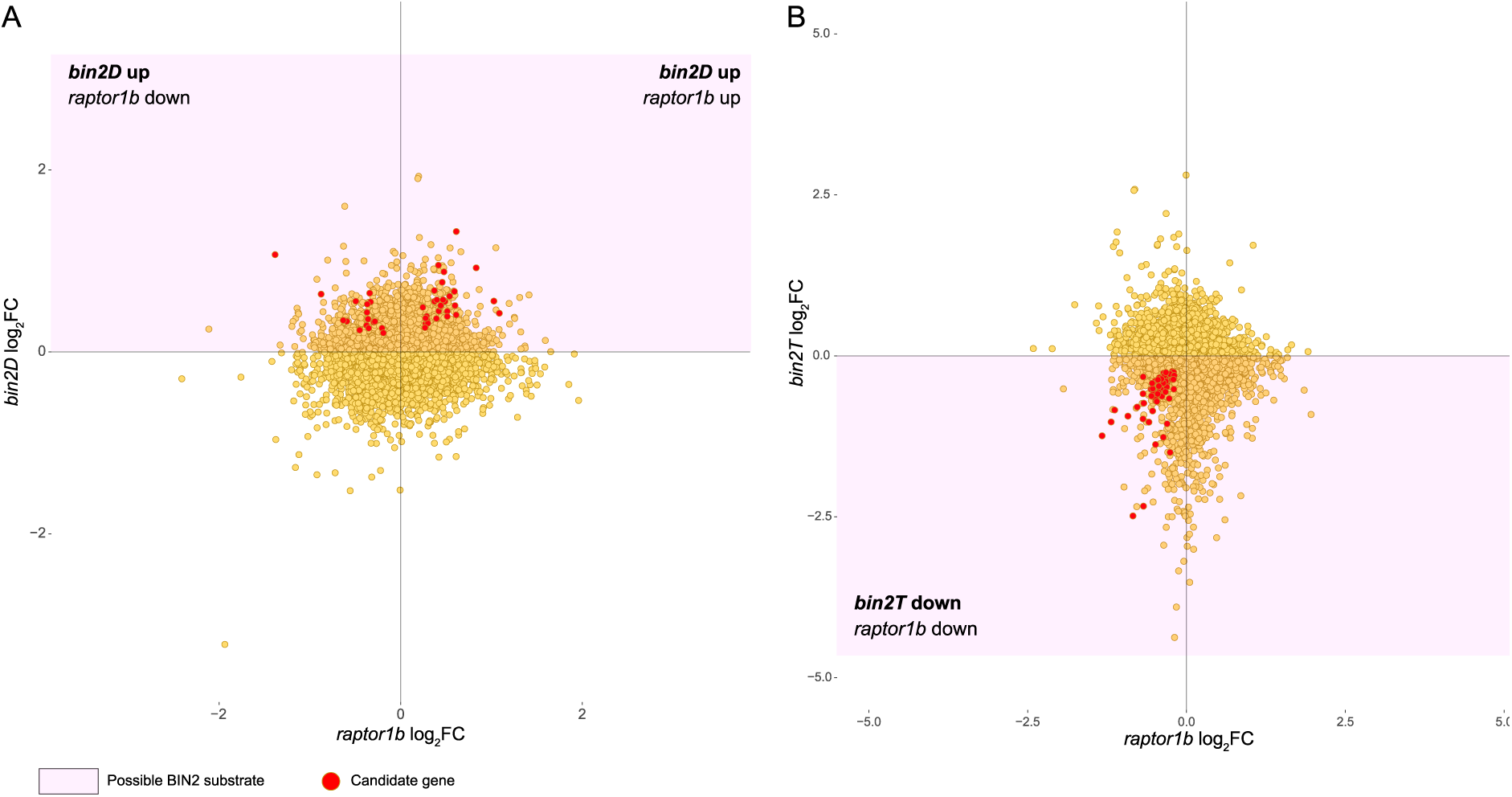
Phosphosite levels in *bin2D, bin2T*, and *raptor1b*. Scatterplot showing phosphosite intensity log2 fold-change (mutant/WT) for *raptor1b* on the x-axis and *bin2D* (**A**) or *bin2T* (**B**) on the y-axis. Red dots indicate phosphosites from selected candidate genes

**Figure 7.**
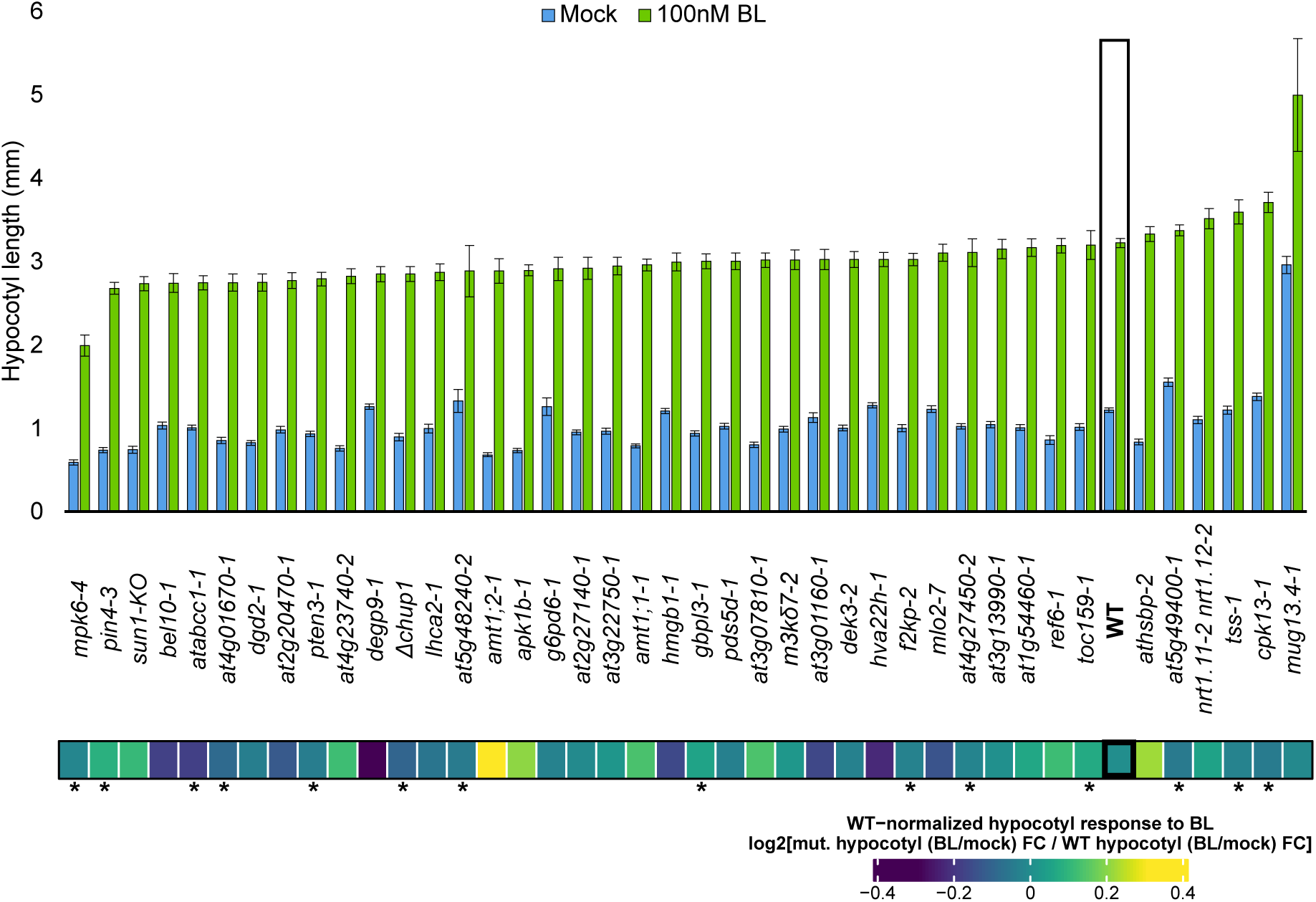
Hypocotyl response to BL treatment in selected mutant lines. (Top) Average hypocotyl length on assayed mutants upon BL treatment. Bar plot showing average (n=24) hypocotyl length on mock (blue bars) or BL (green bars). Wild-type hypocotyl measurement in figure is the average across all experiments and is used only as visual reference. Error bars show standard error. (Bottom) Heatmap showing hypocotyl length response to BL treatment. Values shown are the log2 fold change in hypocotyl length (BL/Mock); n=24. * indicates *p* < 0.1 was observed in each of two independent experiments (Supplemental Dataset S11A).

**Figure 8.**
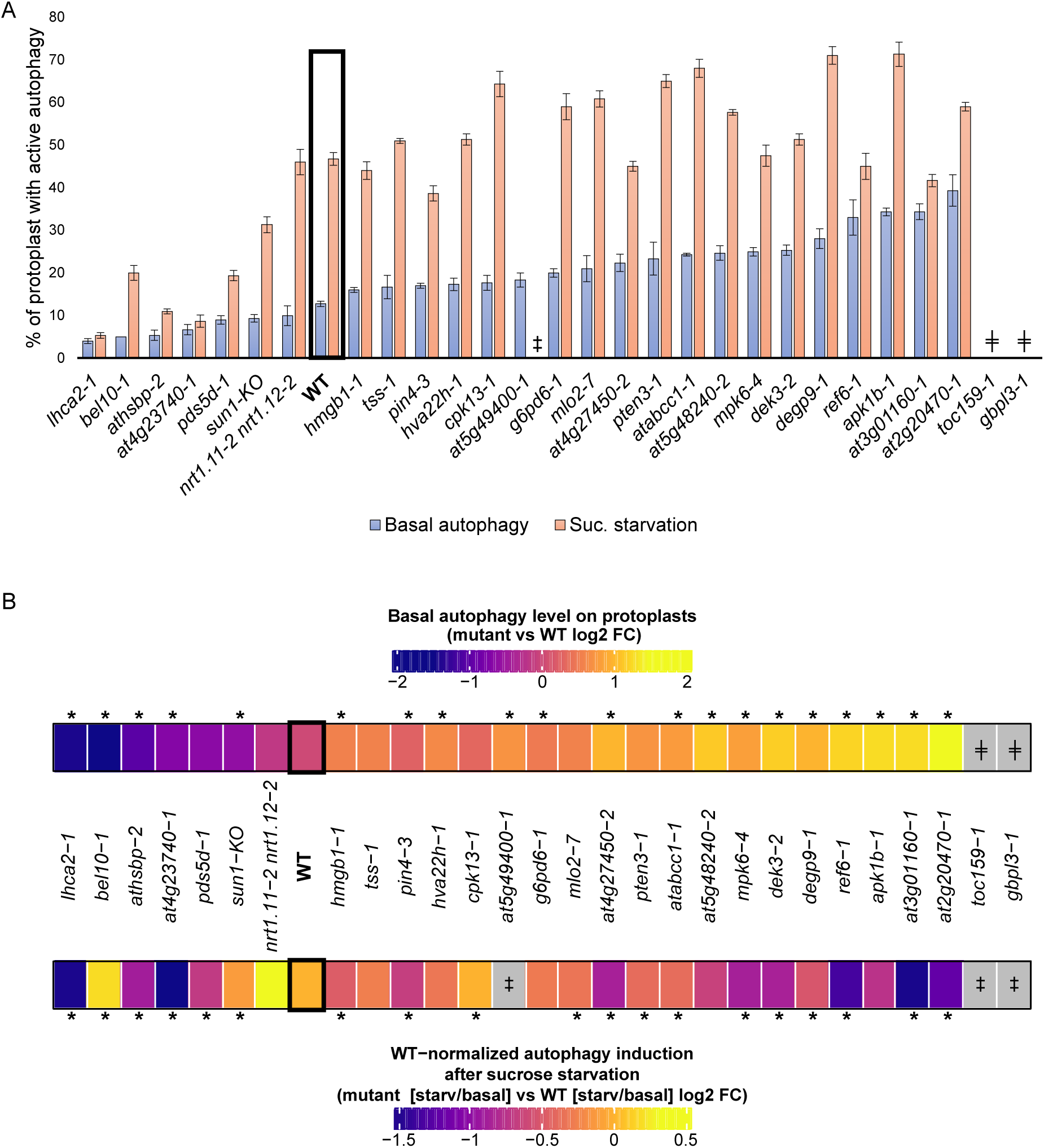
Autophagy levels on selected mutants. **A.** Percentage of protoplast with high autophagy activity under normal conditions (basal autophagy, blue bars) and upon sucrose starvation (orange bars). **B.** Heatmap showing fold-change in protoplasts with active autophagy between mutants and WT protoplasts. (Top) Basal autophagy (mutant/WT) log2 fold-change. * p < 0.05, two-sample t-test. (Bottom) Mutant autophagy response to sucrose starvation (starvation/mock) versus WT response to sucrose starvation (starvation/mock). Values are log2 fold-change. * p < 0.05, generalized linear model. For all measurements, one-hundred protoplast were assessed in triplicate. Error bars show standard error. (╪) Cells from these genotypes did not survive protoplasting. (‡) Cells from these genotypes did not survive sucrose starvation.

In summary, we found a total of 26 genes out of the 41 selected candidates (63.4%) with an altered response to BR and/or level of autophagy. These results confirm the robustness of our integrative multi-omics approach as a way of selecting candidate proteins related to the brassinosteroid and/or autophagy pathways.

## Discussion

Brassinosteroid and TORC have emerged as two key signaling pathways coordinating growth and stress responses. To gain a global view of BR and TORC dependent signaling, we performed deep multi-omics profiling of the transcriptome, proteome, and phosphoproteome of *bin2* and *raptor1b* mutants. These results provide a robust system-wide roadmap of molecular events for BR and TORC pathways crosstalk. Using the multi-omics data, we generated a kinase-signaling network, two TF-centric GRNs, and merged them into one integrative multi-dimensional network, which predicted important genes functioning in BR/TORC pathways. Our genetic studies indeed confirmed that many of the genes predicted from the network play important roles in BR-regulated growth and/or autophagy.

Our work supports previous transcriptome profiling studies of BR including (Wang et al., 2014; Kim et al., 2019; Liu et al., 2020) and TOR signaling (Ren et al., 2012; Xiong et al., 2013; Dong et al., 2015). Despite long-standing interest in BRs, comprehensive (phospho)proteomic profiles examining BR signaling are limited and proteome-wide identification of substrates of the key regulatory kinase BIN2 is lacking. By profiling *bin2* mutants, we identified transcripts, proteins, and phosphorylation sites whose proper expression is dependent on BIN2. Furthermore, using MAKS we provide a global catalogue of potential direct BIN2 substrates. In terms of TORC signaling, we substantially expand on the work of Salem et al., which provided an initial description of proteins that are mis-expressed in *raptor1b* (Salem et al., 2018) as well as the proteins and phosphorylation sites that respond to TOR inhibition via treatment with Torin 2, AZD8055, or rapamycin (Van Leene et al., 2019; Scarpin et al., 2020). Most importantly, through the generation and analysis of these multi-omics data we found a large overlap of gene-products (i.e., transcript, protein, or phosphosites) that are regulated in response to the misexpression of both BIN2 and RAPTOR1B. Together our data suggest extensive crosstalk between BR- and TORC-dependent signaling pathways.

Using our multi-omics data, we reconstructed an integrated gene regulatory and kinase-signaling network. By focusing on the 1,044 regulators and targets predicted by this regulatory network to be co-regulated by BIN2 and RAPTOR1B (Figure 5, center), and accounting for the phosphosites fold-change “directionality” on each mutant (Figure 6), we identified and tested a set of 41 candidate genes for their involvement in BR/TORC signaling pathways. Mutants for these 41 genes were assayed and 26 of them showed an altered phenotype.

To summarize these phenotyping results, we divided our gene set into groups according to their different phenotypes in BR-regulated hypocotyl elongation and autophagy levels as a readout of TORC activity (Figure 9). TOR signaling is known to be positively regulated by auxin and glucose availability (Xiong et al., 2013; Li et al., 2017; Schepetilnikov et al., 2017). Here, we found that mutants in the auxin efflux carrier PIN4 exhibited increased autophagy (Figure 9, purple). This finding agrees with their corresponding functions: PIN4 creates an auxin sink at the cells below the quiescent center, a crucial event for normal auxin distribution (Friml et al., 2002). In parallel with this, we discovered homologs of proteins involved in autophagy and TOR signaling in human. *Homo sapiens* (Hsa) PTEN has been shown to negatively regulate both mTOR signaling and autophagy through independent pathways (Errafiy et al., 2013). In agreement, our results show that Arabidopsis PTEN3 mutant plants have increased autophagy (Figure 9, purple). Conversely, HsaHMGB1 can translocate to the cytoplasm and induce autophagy upon perception of reactive oxygen species (Tang et al., 2010). Here, HMGB1 mutants show reduced sensitivity to BR and increased autophagy levels (Figure 9, dark green), suggesting an opposite function in plants.

**Figure 9.**
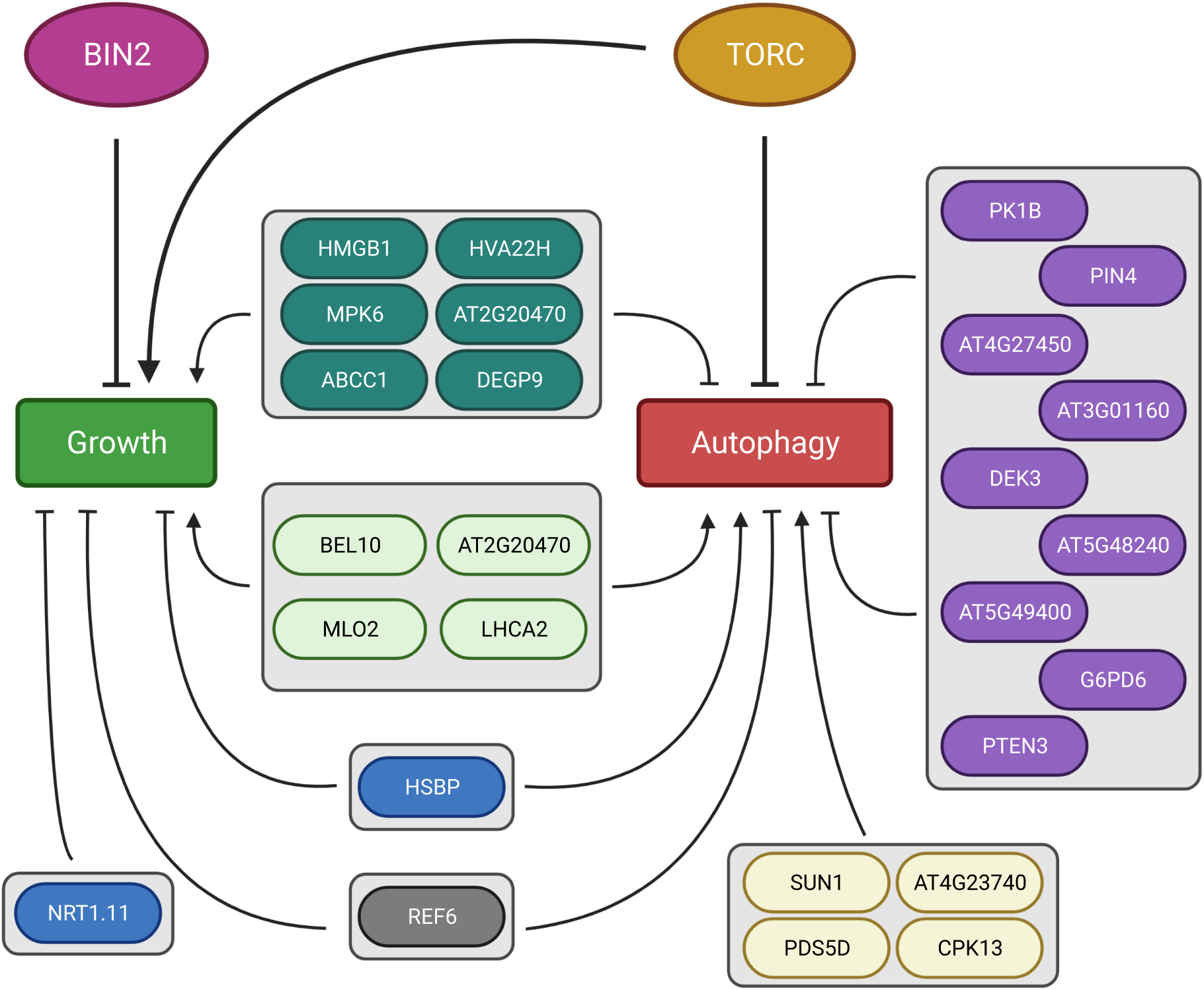
Proposed model of interaction for significant genes. Genes with significant response to BL or altered autophagy levels are organized into groups according to their mutants’ phenotype.

Finally, our results identified previously unknown genes involved in BR and autophagy pathways. For example, mutant phenotype analysis indicated that plants with mutated MPK6 show reduced sensitivity to BR and increased autophagy levels (Figure 9, dark green). Although no direct role as a BR-induced growth has been established, MPK6 kinase is involved in a myriad of processes and has been shown to directly phosphorylate and activate BES1 to increase immune response (Kang et al., 2015). Furthermore, BIN2 can phosphorylate and inhibit MKK4, a direct MPK6 activator (Khan et al., 2013). Moreover, our signaling network prediction situates MPK6 as potentially being upstream of RAPTOR1B, which will be interesting to test in future work. Many other genes that may play a role in both BR-regulated growth and modulation of autophagy were identified by our analysis (Figure 9).

In summary, this study builds upon previous findings that BR and TORC crosstalk in the regulation of plant growth and stress responses (Zhang et al., 2016; Nolan et al., 2017b; Xiong et al., 2017; Vleesschauwer et al., 2018; Liao and Bassham, 2020; Nolan et al., 2020). Our multi-omics studies provide genome-wide evidence for extensive interactions between BR and TORC signaling pathways across three different regulatory layers (i.e., transcript, protein, and phosphorylation). These results establish an integrative signaling network that defines molecular interactions between BR-or TORC-regulated growth and autophagy.

## Methods

### Plant material

Arabidopsis thaliana mutant lines *bin2-1* (Li and Nam, 2002), *bin2-3 bil1 bil2* (Yan et al., 2009), and *raptor1-1* (Anderson et al., 2005; Deprost et al., 2005) were used in this research as *bin2D, bin2T*, and *raptor1b*, respectively. The full list of seed stocks used in this work are summarized in Supplemental Data Set S12. All plants were grown in LC1 soil mix (Sungro) under long day conditions (16 h light, 22°C) unless stated otherwise. Columbia-0 ecotype was used as wild-type control for all assays.

### QuantSeq library preparation and sequencing

Four biological replicates of 20-day-old rosette leaves were collected from WT and each mutant (*bin2T, bin2D*, and *raptor1b*) and immediately frozen in liquid N2. Tissue was ground for at least 15 minutes under liquid N2 using mortar and pestle. Total RNA was extracted using RNAeasy Plant Mini Kit with DNaseI treatment (Qiagen). Five-hundred ng of total RNA was used for QuantSeq 3’ mRNA-seq library Prep kit FWD for Illumina (Lexogen) (Moll et al., 2014). Library sequencing was performed on an Illumina HiSeq3000.

### Transcriptomic data analysis

Data processing followed the pipeline suggested by QuantSeq 3’ mRNA-seq Integrated Data Analysis Pipelines on Bluebee® Genomic Platform User Guide (Lexogen Cat. 090-094). Briefly, reads were adapter- and quality-trimmed using BBDuk v37.36. Trimmed reads were mapped to the Arabidopsis reference transcriptome (TAIR10 annotation) using Star Aligner v2.5.3a (Dobin et al., 2013). Finally, transcript counts were extracted using HTSeq-count v0.11.2 (Anders et al., 2015).

Differential expression was assessed using the PoissonSeq R package (Li et al., 2012). A q-value < 0.05 and fold change > 1.3 (log2FC > 0.4) was used as cutoff for designating differentially expressed transcripts. All data processing scripts were deposited in a github repository (see Data Availability section)

### Protein extraction for global proteome and phosphoproteome profiling

Three biological replicates from the same tissue collected for transcriptome analysis were used for protein extraction via the urea-FASP method described by (Song et al., 2018a, 2018b, 2020) as follows: 250 mg of finely ground tissue was subjected to mechanical disruption using 250 mg ZrO2 beads in presence of 500 µL lysis buffer (8M urea; 100 mM TRIS-HCl, pH 7.0; 5mM TCEP) on a MiniG tissue homogenizer (Spex SamplePrep). Sample was clarified, and supernatant was transferred to a clean tube. Proteins were precipitated using rounds of 45-minute incubation at -80°C as follows: 1 round of ice-cold 100% acetone, 2 rounds of 80% acetone and 3 rounds of 0.2mM Na3VO4 in 100% methanol. Each round of incubation consisted of sample resuspension assisted by probe sonication, incubation at -20°C, centrifugation at 4,500xG for 10 minutes, and removal of supernatant. Extracted proteins were resuspended by sonication in urea resuspension buffer (URB, 8M urea in 50 mM TRIS-HCl, pH = 7.0; 5 mM TCEP; 1x phosphatase inhibitor cocktail) and further cleaned through Filter-Assisted Sample Preparation (FASP) using Amicon Ultra-4 30kDa MWCO filter units (Millipore) in the presence of UA buffer (8M urea in 100mM TRIS-HCl, pH = 8.0; 1x phosphatase inhibitor cocktail). Samples were reduced with 2mM TCEP, alkylated using 50 mM iodoacetamide (IAM) and digested into peptides using one round of overnight incubation at 37°C with 1:50 (enzyme:protein) trypsin (Roche, Cat. No. 03708969001) and a second round of incubation for 4 hours at 37°C with trypsin and Lys-C (Wako Chemicals, Catalog number 125-05061). Purified samples were further desalted using SepPack C18 columns (Waters). Tandem Mass Tag (TMT lot #TC264166, Thermo Scientific) labeling was performed on 330 µg of purified peptides from each sample in a 1:1.7 (peptide:label) ratio as previously reported (Song et al., 2020). TMT labeling reaction efficiency was assessed to be at least of 98% by LC-MS. Following verifying labeling efficiency the reaction was quenched using 5% hydroxylamine and the samples were then pooled. One hundred µg of labeled peptide was set aside for global proteome profiling and the remaining labeled sample was subjected to a second round of C18 desalting before phosphopeptide enrichment. Serial Metal Oxide Affinity Chromatography (SMOAC) method from Thermo Scientific was used for phosphopeptide enrichment. Briefly, High-Select TiO2 Phosphopeptide Enrichment kit (Thermo Scientific) was used as a first enrichment step and all flow-through was pooled, concentrated to almost dry on a SpeedVac and used as input for High Select Fe-NTA Phosphoptide Enrichment kit (Thermo Scientific). Phosphopeptides obtained from both enrichment processes were dried using a SpeedVac, resuspended in 0.1% formic acid in Optima grade H2O (Millipore) and pooled together.

### BIN2 Multiplexed Assay for Kinase Specificity

MAKS was performed based on the protocol described by (Jayaraman et al., 2017). Protein was extracted for MAKS from 1 g of leaf ground tissue from 20-days-old wild type Arabidopsis plants using the phenol-FASP protocol as described previously (Song et al., 2018b, 2020). Three mg of total purified protein was resuspended in URB, re-precipitated in ice-cold 100 mM NH4CH3CO2 in 100% methanol. Following precipitation, the solvent was removed and the protein pellet was resuspended in kinase buffer (50 mM TRIS-HCl, pH = 7.7; 5 mM MgCl2; 5mM ATP; 1x phosphatase inhibitor cocktail). Resuspended protein was divided into 600 µg aliquots and incubated with either recombinant GST or GST-BIN2 at a 1:75 (enzyme:protein) ratio at 37°C with gentle shaking for 1 hour. After incubation, protein solution was subjected to FASP, reduced with 2mM TCEP, alkylated in 50mM IAM, and digested using trypsin as described by (Song et al., 2020). Three replicates were made for each treatment (i.e., GST and GST-BIN2). Two hundred µg of peptides from each replicate were used for TMT labeling. Phosphopeptide enrichment was performed on labeled peptides using SMOAC as described previously in this paper. Cloning of GST-BIN2 was described in (Yin et al., 2002); The fusion protein was purified using Glutathione agarose beads as described in (Jiang et al., 2019).

### LC-MS/MS

Chromatography was performed on an Agilent 1260 quaternary HPLC with constant flow rate of ∼600 nL min^-1^ achieved via a splitter. A Sutter P-2000 laser puller was used to generate sharp nanospray tips from 200 µm ID fused silica capillary. Columns were all in-house packed on a Next Advance pressure cell using 200 µm ID capillary. All samples were loaded into a 10 cm capillary column packed with 5 μM Zorbax SB-C18 (Agilent) and then connected to a 5 cm-long strong cationic exchange (SCX) column packed with 5µM PolySulfoethyl. The SCX column was then connected to a 20 cm long nanospray tip, packed with 2.5 µm C18 (Waters). For global protein abundance, 45 µg of labeled peptides were fractionated online using 27 ammonium acetate salt steps. For phosphoproteomics, 25 µg of enriched peptides and 14 salt steps were used. For MAKS, 30 µg of enriched peptides and 17 salt steps were used. Each salt step was then separated using a 150 min reverse-phase gradient (Zhang et al., 2019).

Eluted peptides were analyzed using a Thermo Scientific Q-Exactive Plus high-resolution quadrupole Orbitrap mass spectrometer, which was directly coupled to the HPLC. Data dependent acquisition was obtained using Xcalibur 4.0 software in positive ion mode with a spray voltage of 2.1 kV and a capillary temperature of 275 °C and an RF of 60. MS1 spectra were measured at a resolution of 70,000, an automatic gain control (AGC) of 3e6 with a maximum ion time of 100 ms and a mass range of 400-2000 m/z. Up to 15 MS2 were triggered at a resolution of 35,000. A fixed first mass of 115 m/z. An AGC of 1e5 with a maximum ion time of 50 ms, a normalized collision energy of 33, and an isolation window of 1.3 m/z for global proteome and 1.5 m/z for phosphoproteome were used. Charge exclusion was set to unassigned, 1, 5–8, and >8. MS1 that triggered MS2 scans were dynamically excluded for 25 s for global proteome and 45 s for phosphoproteome.

### Proteomics data analysis

Spectra for global protein abundance runs were searched using the Andromeda Search Engine (Cox et al., 2011) against the TAIR10 Arabidopsis proteome (https://www.arabidopsis.org/download_files/Proteins/TAIR10_protein_lists/TAIR10_pep_20101214) using MaxQuant software v1.6.1.0 (Tyanova et al., 2016). Carbamidomethyl cysteine was set as a fixed modification while methionine oxidation and protein N-terminal acetylation were set as variable modifications. Digestion parameters were set to “specific” and “Trypsin/P;LysC”. Up to two missed cleavages were allowed. A false discovery rate less than 0.01 at both the peptide spectral match and protein identification level was required. Sample loading and internal reference scaling normalization methods were used to account for differences within and between LC-MS/MS runs, respectively (Plubell et al., 2017).

Differential expression was assessed using the PoissonSeq R package (Li et al., 2012). A q-value < 0.1 was used as cutoff for designating differentially expressed proteins. Scripts for data analysis were deposited in a github repository (see Data Availability section)

### Phosphoproteomics data analysis

Spectra for both *bin2*/*raptor1b* mutant profiling and MAKS were searched together using similar approach as with global protein abundance with exceptions. Briefly, MaxQuant software v1.6.10.43 was used instead and “Phospho (STY)” search for variable modifications was included. Sample loading and internal reference scaling normalization methods were used to account for differences within and between LC-MS/MS runs, respectively (Plubell et al., 2017).

Differential expression was assessed using the edgeR R package (Robinson et al., 2010). A q-value < 0.1 was used as cutoff for designating differential phosphorylation. See Data Availability section for the full analysis script.

### Motif enrichment analysis

Motif enrichment was performed using the motifeR R package (Wang et al., 2019) with default settings: serine or threonine as the central residues, a p-value threshold of 0.001, and TAIR10 protein annotation as background reference. Enrichment p-value was calculated using hypergeometric testing using *phyper* function in R.

### Analysis of overlap between BIN2 MAKS and *bin2* mutant datasets

To find overlapping phosphosites, we defined any two distinct phosphosites as identical if they originated from the same phosphoprotein and were less than 10 amino acid residues apart. This approach was used to account for cases where phosphosites were not localized to a specific amino acid on a given peptide in the two different datasets. Overlap statistical significance was assessed by hypergeometric test.

### Kinase activation loop prediction

Protein kinases were identified using a modified version of the pipeline described by (Walley et al., 2013; Clark et al., 2021). Briefly, all 35,386 protein sequences available in the TAIR10 annotation (https://www.arabidopsis.org/download_files/Proteins/TAIR10_protein_lists/TAIR10_pep_20101214) were searched for kinase domain using The National Center for Biotechnology Information batch conserved domain search tool (Lu et al., 2020). From this list of 1,522 proteins with identified kinase domain, 878 were also annotated with activation loop (A-loop) coordinates by the search tool. The kinase domains of proteins lacking the A-loop coordinates were aligned using MAFFT (Katoh and Standley, 2013) and the well conserved A-loop beginning (DFG) and end (APE) motifs were manually searched. An extra 482 A-loop coordinates were obtained, for a total of 1,360 protein kinases with A-loop coordinates.

### Kinase-signaling network

Kinases with differential phosphorylation inside the A-loop (activated kinases) were used as regulators to build the kinase-substrate network. For this, the Spearman and Pearson correlation between a regulator and the rest of differentially phosphorylated peptides was calculated as described previously (Supplemental Figure 2B) (Walley et al., 2013; Clark et al., 2021).

### Gene Regulatory Network reconstruction

A curated list of transcriptional regulators was used to identify quantified TFs in our datasets (Supplemental data set S13) For the abundance GRN, TF protein abundance (when quantified) or TF transcript abundance (when cognate protein was not quantified) was used as the “regulator” value to infer their “target” transcript abundance. The phosphosite GRN, uses the TF phosphorylation intensity value as “regulator” to predict “target” transcript abundance. In order to mix different data sources (i.e., proteomics, phosphoproteomics, and transcriptomics) into consolidated tables, expression values for each “omics” were rank-normalized using the *norm*.*rrank* function from the r package “demi” (Ilmjärv et al., 2014).

In both networks, transcript abundance was used to build the target tables.

Network inference was achieved using a modified version of the GENIE3 random forest algorithm (Huynh-Thu et al., 2010) in the SC-ION pipeline V2.1 (https://doi.org/10.5281/zenodo.5237310) with no clustering, as described before (Clark et al., 2021). Results were visualized in Cytoscape v3.9.0 (Shannon et al., 2003) IVI score was calculated using the “influential” and “igraph” r packages following developer instructions (Csardi and Nepusz, 2006; Salavaty et al., 2020). A table with all the network interactions, where the first column had the regulator’s gene ID and the second column had the corresponding target’s gene ID, was parsed using *graph_from_data_frame* function and graph vertices were obtained using *GraphVertices* function. Before calculating IVI score, the following centrality measurements were needed and calculated: degree centrality, betweenness centrality, neighborhood connectivity, collective influence, clusterRank, and local H-index. IVI was calculated using the function *ivi*.*from*.*indices*.

### BL response assays

Seeds were vapor-phase sterilized in chlorine gas, stratified at 4°C for 1 week, and germinated in on petri plates containing half-strength Linsmaier and Skoog media (Caisson Labs, catalog number LSP03) in 0.7% Phytoblend (Caisson Labs, catalog number PTP01), supplemented with 1% sucrose and either DMSO or 100 nM brassinolide (BL). Seedlings were grown for 7 days at 22/18°C (day/night), 16 hours of light, 40% relative humidity, and light intensity of 120 µmol m^-2^ s^-1^. Seedlings were imaged and hypocotyl length was measured using Fiji software (Schindelin et al., 2012). Twenty-four seedlings per mutant were used on each treatment and this experiment was repeated at least two times for those genotypes showing significant response. A generalized linear model with treatment and genotype as factors and controlling for random effects of replicate and plate was applied using the *glmmPQL* function from the MASS R package (Venables and Ripley, 2002), and a threshold of “genotype by treatment interaction”. A p-value < 0.1 in two independent experiments was set as the significance cutoff.

### GFP-ATG8e protoplast assay

Protoplasts were extracted from leaves from 20-day-old plants and transformed as described previously (Wu et al., 2009). Protoplasts were observed by epifluorescence microscopy (Carl Zeiss Axio Imager.A2, Germany) using a FITC filter, and protoplasts with more than three visible autophagosomes were counted as active for autophagy as previously described (Yang et al., 2016; Pu et al., 2017). One hundred protoplasts were analyzed per treatment per genotype and the experiment was repeated three times. For sucrose starvation, transformed protoplasts were incubated in W5 solution without sucrose or with 0.5% (w/v) sucrose as control at room temperature for 36 hours in dark before assessing autophagy.

Significance of basal autophagy levels was assessed by two-sample t-test whereas a generalized linear model with treatment and genotype as factors and controlling for random effects of replicates was used for autophagy levels under sucrose starvation. A p-value < 0.05 was used as a cutoff on both cases.

## Acknowledgements

This work was supported by the Iowa State University Plant Science Institute (YY and JW), NIH R01GM120316 (YY, DB, JWW), NSF IOS-1818160 (YY and JW), and USDA NIFA Hatch project IOW3808 funds to JWW. NMC is supported by a USDA NIFA Postdoctoral Research Fellowship (2019-67012-29712) and TMN is supported by the National Science Foundation Postdoctoral Research Fellowships in Biology Program (Grant No. IOS-2010686). We thank Peng Liu (ISU Department of Statics) for help in determining statical analysis for the BL and autophagy phenotype assays.

## Figure Legends

**Supplemental Figure S1. Differential expression analysis**. UpSet plot showing overlap of DE transcripts (**A**), proteins (**B**), and phosphosites (**C**) between *bin2D, bin2T*, and *raptor1b* mutants.

**Supplemental Figure S2. Workflow of signaling network reconstruction. A**, Pipeline for activation loop coordinates annotation for Arabidopsis kinases. **B**, Logic diagram for kinase/substrate prediction.

