## Supplementary figures and images for "Integration of multi-omics data reveals interplay between brassinosteroid and TORC signaling in Arabidopsis"

### Supplemental Figure S1

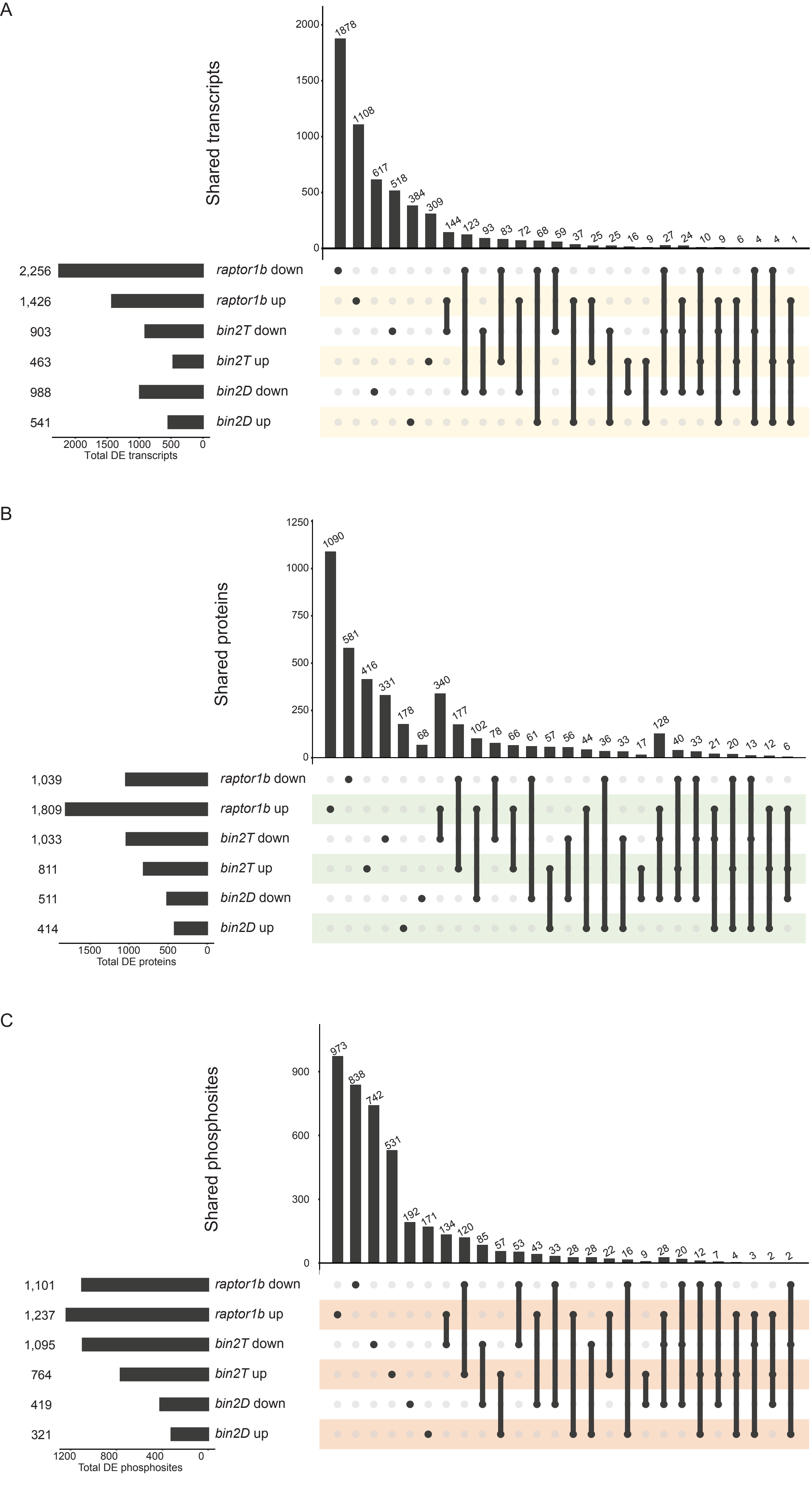

### Supplemental Figure S2

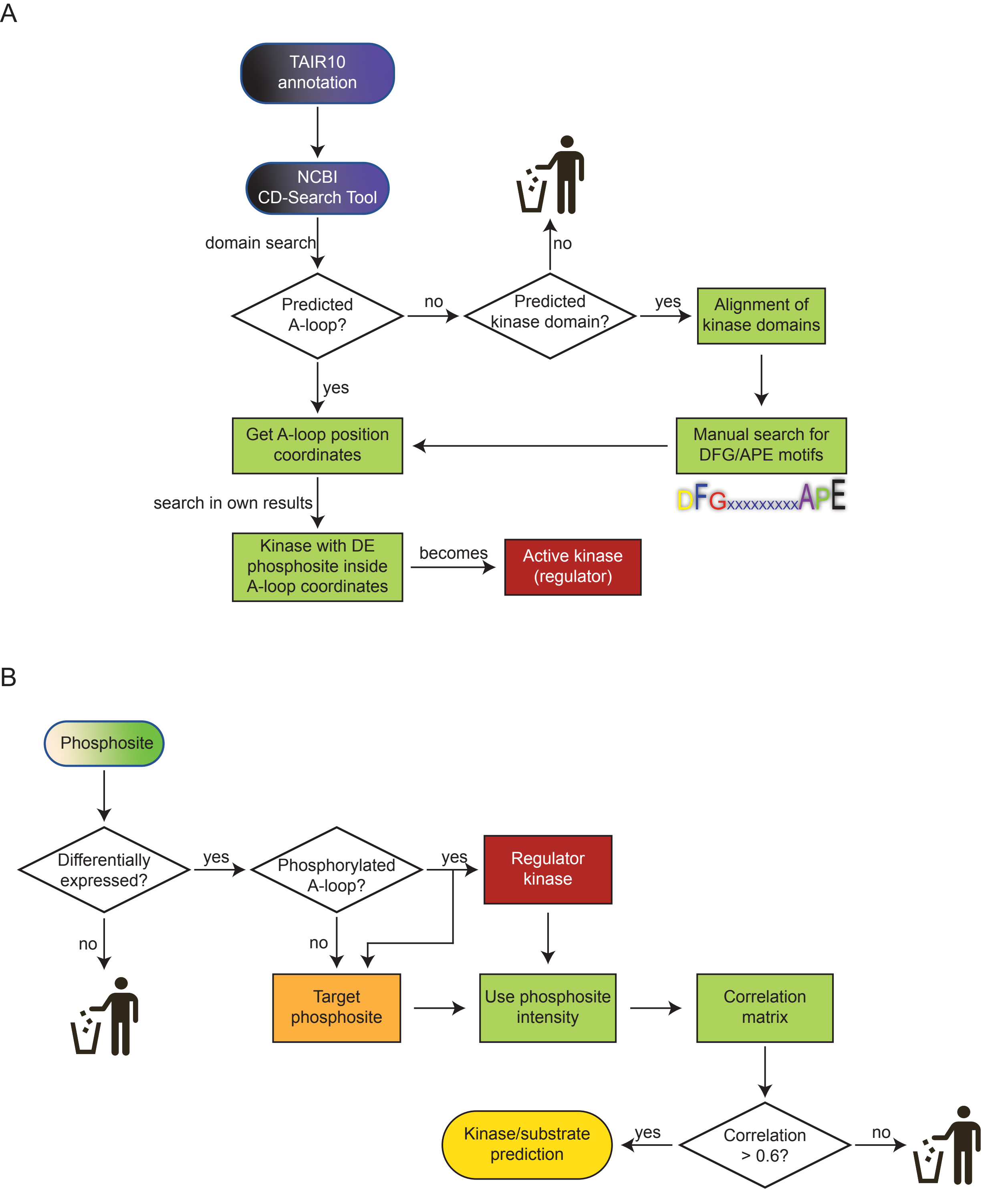


Supplemental Figure S2
